# A stress-activated mid-insula to BNST pathway regulates susceptibility to abstinence-induced negative affect in female mice

**DOI:** 10.1101/2025.01.07.631325

**Authors:** Benjamin M. Williams, Jincy R. Little, Nathaniel S. O’Connell, Samuel W. Centanni

**Affiliations:** Department of Translational Neuroscience, Wake Forest University School of Medicine, Winston-Salem, NC, USA; Department of Biostatistics and Data Science, Wake Forest University School of Medicine, Winston-Salem, NC, USA

## Abstract

Stress is central to many neuropsychiatric conditions, including alcohol use disorder (AUD). Stress influences the initiation and continued use of alcohol, the progression to AUD, and relapse. Identifying the neurocircuits activated during stress, and individual variability in these responses is critical for developing new treatment targets for AUD, particularly to mitigate stress-induced relapse. Using a longitudinal approach, this study examined the relationship between sub-chronic stress exposure and negative affect during protracted abstinence following chronic ethanol exposure. Sub-chronic restraint stress heightened negative affect-like behavior in protracted abstinence. Interestingly, this was driven by a subset of “stress-susceptible” female mice. We examined the mid-insula, a hub in the brain’s salience network, as a driver of this effect, given its role in emotional regulation and links to alcohol craving, consumption, and abstinence-induced negative affect. Mid-insula GCaMP fiber photometry revealed that GCaMP activity during stress exposure was positively correlated with activity during the novelty-suppressed feeding test (NSFT) two weeks into abstinence. A distinct subset of mice exhibited increasing activity during the consummatory phase, implicating the mid-insula as a neural basis for heightened negative affect in abstinence. Chemogenetic inhibition of mid-insula neurons projecting to the dorsal BNST during stress disrupted the emergence of stress susceptibility, highlighting this circuit as a key determinant of susceptibility to abstinence-induced negative affect. These outcomes were female-specific, addressing a critical gap in understanding AUD risk in women. Furthermore, female mice exhibited higher struggling behavior during stress than males. However, this effect was blocked by chemogenetic inhibition of the insula-BNST pathway during stress. By linking pre-alcohol stress response with abstinence outcomes, this work positions the insula-BNST pathway as a potential AUD circuit activity biomarker and therapeutic target.

## INTRODUCTION

Stress is a pervasive and individualized experience that significantly contributes to the onset and exacerbation of numerous neuropsychiatric diseases, including mood, substance use, and anxiety disorders. Among these, alcohol use disorder (AUD) stands out due to its profound impact on public health. Stress and negative emotional states are intertwined with the initiation and continuation of alcohol use, and, in some individuals, the progression to alcohol use disorder (AUD) [1]. AUD patients in abstinence often experience hyperkatifeia, or heightened sensitivity to emotional distress in drug or alcohol withdrawal [2]. The inability to properly cope with stress is a key driver of relapse and cravings in many individuals [3]. Despite this link, using stress response as a predictive marker of disease manifestation and progression has been unsuccessful. A primary challenge lies in the complexity of a stress response. Stressors can have drastically different short- and long-term impacts on individuals, and mental health conditions involve complex overlapping dysfunctional patterns that complicate pinpointing cause and effect. Furthermore, sex differences in the progression to alcohol dependence and susceptibility to negative affect-related disorders are well-established [4–8], yet few models have encapsulated this, limiting a deeper understanding of the neurobiological mechanisms driving these differences.

Neurocircuits are the basic functional units that encode behavior and could serve as a more effective indicator of AUD-related behavior like susceptibility to hyperkatifeia and stress-induced relapse, thus informing more individualized treatment strategies. The insula, a central hub in the brain’s salience network, regulates the switch between the default mode network to the executive control network [9,10] and conveys interoceptive cues through cortical and subcortical connections to predict outcomes and guide behavioral actions [11–13]. Insula integration of stimuli occurs in a posterior-to-anterior gradient, with initial salience detected in the posterior insula and higher cognitive processing in the anterior insula [14]. Positioned between these subregions, the mid-insula is considered critical for emotional integration. Abnormalities in insula function have been implicated in mood disorders and AUD [11], with human imaging studies showing increased insula activation in response to alcohol cues [15,16], psychosocial stress, and uncertain threats [17,18]. Our previous work established a role for the mid-insula in alcohol abstinence-induced negative affect and stress response [19,20]. These findings underscore the insula’s role in cravings, consumption, and abstinence-related emotional dysregulation.

Despite the vast connectivity of the insula, its specific projections, such as those to the extended amygdala [19,21], have received comparatively little attention. The dorsal bed nucleus of the stria terminalis (BNST), a component of the extended amygdala that receives dense insula projections, integrates negative valance or affective states driven by cortical, subcortical, midbrain, and hindbrain inputs [22,23]. Human fMRI studies have validated a structural and functional insula and BNST connection [24,25]. Preclinical studies from our lab and others defined a role for the insula to ventral BNST circuit in binge alcohol drinking [26] and the dorsal BNST in negative affect-like behavior in protracted alcohol abstinence [19,20,26] and active coping behavior during restraint stress [20], making for a compelling candidate to explore susceptibility to AUD-related behavior like hyperkatifeia in alcohol abstinence.

This study builds off the independently defined roles of the insula-BNST pathway in stress response [20] and alcohol abstinence [19]. Using a longitudinal mouse model that combines sub-chronic restraint stress, continuous access two-bottle choice alcohol drinking, and protracted abstinence, we sought to link pre-alcohol stress response with drinking patterns and negative affect in abstinence. We employed in vivo calcium imaging and circuit-specific chemogenetics to dissect the functional contributions of the insula→BNST pathway. This work provides a mechanistic framework linking stress-induced neuronal activity to AUD-related behaviors. By identifying the mid-insula-BNST pathway as a key modulator of stress-susceptibility, this study offers novel insights into the neurobiological underpinnings of AUD vulnerability, particularly abstinence-induced negative affect. These findings highlight the potential of targeting this pathway for earlier intervention and personalized treatment strategies, shifting the focus from reactive to preventive approaches.

## METHODS AND MATERIALS

### See supplementary material for detailed methods

Singly-housed male and female C57BL/6J mice (n=124; The Jackson Laboratory) were delivered at age seven weeks and acclimated for one week before experimentation. All procedures were conducted with the approval of the Institutional Animal Care and Use Committee at Wake Forest University and were within the guidelines set forth by the Care and Use of Mammals in Neuroscience and Behavioral Research [27].

Sub-chronic restraint stress and CDFA were conducted as previously described [19,20,28] and in Figure 1. All viruses were purchased from Addgene, used as received and stereotaxically injected into the mid-insula and/or the dorsal BNST (300 nL). DREADD agonist 21 (C21, MilliporeSigma) was administered (1 mg/kg, i.p.) 1h before stress. Mice were handled, and behavioral studies were performed as previously described [29–31].

**Figure 1.**
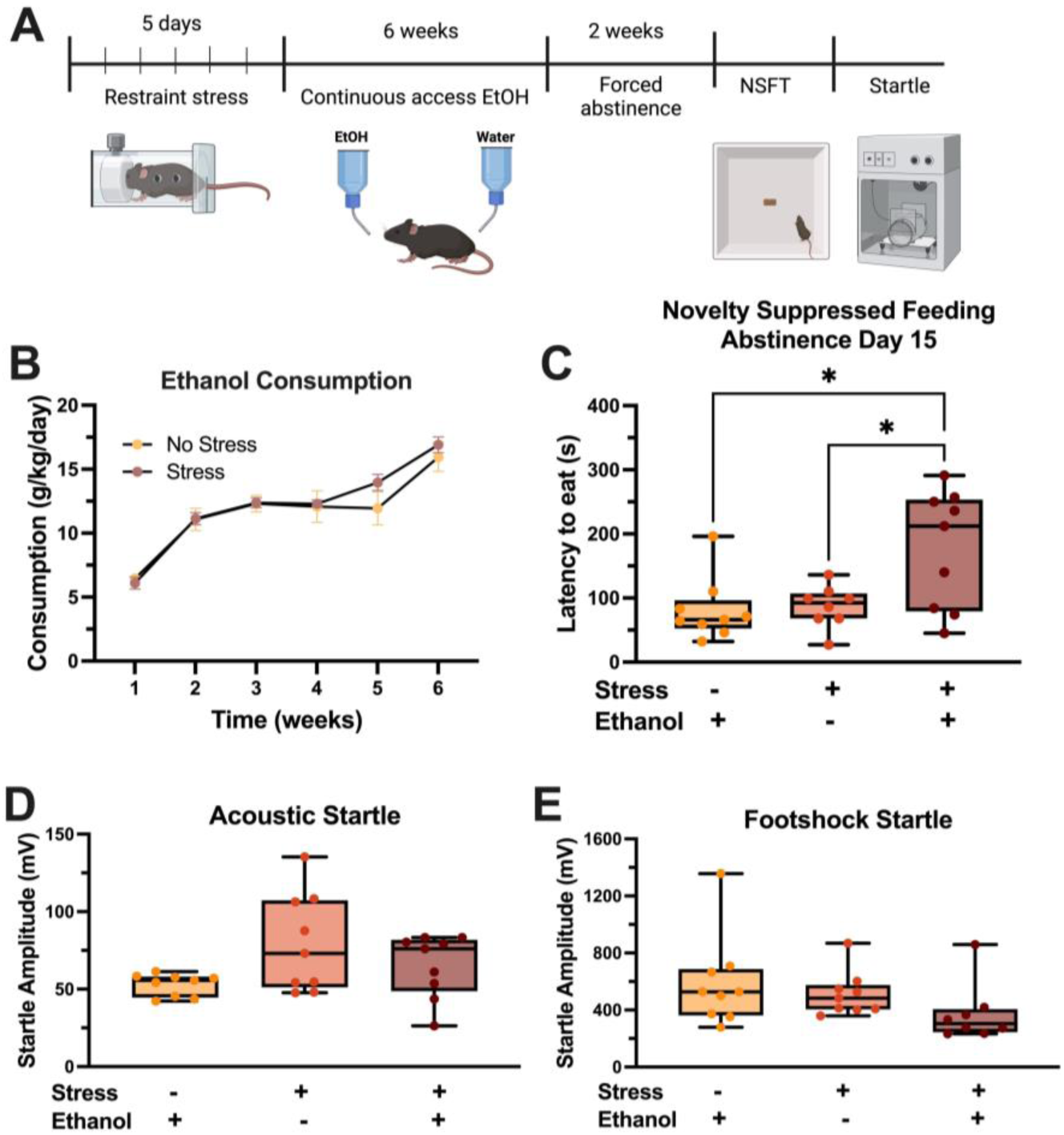
Sub-chronic stress exposure enhanced negative affective phenotypes following 6 weeks of continuous ethanol drinking. (A) Experimental design. (B) No difference in ethanol consumption between stress and no stress groups. (C) Mice with prior stress and alcohol exposure show higher latency to eat the food during NSFT in abstinence, compared to stress and no ethanol controls. (D-E) Acoustic (D) and footshock (E) startle responses are similar between groups. * p<0.05

All data were analyzed using R-Studio and GraphPad Prism 10, with a significance level of p<0.05. For continuous outcomes (e.g., ethanol consumption) across multiple treatment/comparative groups, ANOVA was used with an appropriate method for multiple comparisons (e.g., Dunnet’s test) as relevant to the hypotheses being tested. For simple two-group comparisons, t-tests were used.

## RESULTS

### Sub-chronic restraint stress increases abstinence-induced negative affect-like behavior

To assess the impact of sub-chronic stress on negative affect-like behavior during abstinence, female C57BL/6J mice underwent five days of restraint stress (1h/day) followed by the chronic ethanol drinking-forced abstinence model (CDFA) [19] (Figure 1A). Restraint stress did not impact ethanol consumption (Figure 1B). However, two weeks into abstinence, sub-chronic stress exposure before drinking drove a significantly higher latency to eat during the novelty suppressed feeding test (NSFT) (one-way ANOVA, F_(2,23)_=6.26, p=0.0068, Tukey’s post hoc stress effect p=0.011, ethanol effect p=0.021, Figure 1C), suggesting that stress before CDFA results in higher anxiety-like behavior. Interestingly, this effect was specific to NSFT, as no differences were observed in acoustic and footshock startle responses across groups (Figure 1D-E). Furthermore, a distinct subset of mice drove the increased NSFT latency effect, suggesting stress-susceptible and stress-resilient sub-populations. These sub-populations exhibited no differences in ethanol consumption or startle response (Figure S1), highlighting the specificity of the NSFT finding.

### Mid-insular GCaMP activity during restraint stress is negatively correlated with ethanol consumption

We previously demonstrated a unique role for the mid-insula in stress-coping behavior and ethanol abstinence-induced negative affect [19,32]. We investigated whether mid-insula activity during stress is an indicator of subsequent ethanol consumption. Female mice were injected with GCaMP8f into the mid-insula and implanted with fiberoptics for *in vivo* recordings during restraint stress. Active escape attempts (struggle bouts), a validated measure of negative affect [20,33,34], were quantified using machine learning pose estimation (DeepLabCut) and custom R code (Figure 2A-B). Consistent with prior findings, mid-insula GCaMP activity increased at the onset of a struggle bout (Figure 2C). Peak insula GCaMP response decreased after each day of restraint stress (one-way ANOVA, F_(4,59)_=14.32, p<0.0001, Sidak-Holmes Post hoc p<0.0001 for days 1 vs 3-5, Figure 2D), reflecting desensitization typically observed after repeated homotypic stress exposure. Following stress, mice underwent CDFA. Peak GCaMP amplitude during the first stress was negatively correlated with ethanol consumption during the last three weeks of drinking (R^2^=0.387, F_(1,10)_=6.31, p=0.031, Figure 2E). This suggests a stronger mid-insula response to stress is associated with decreased consumption.

**Figure 2.**
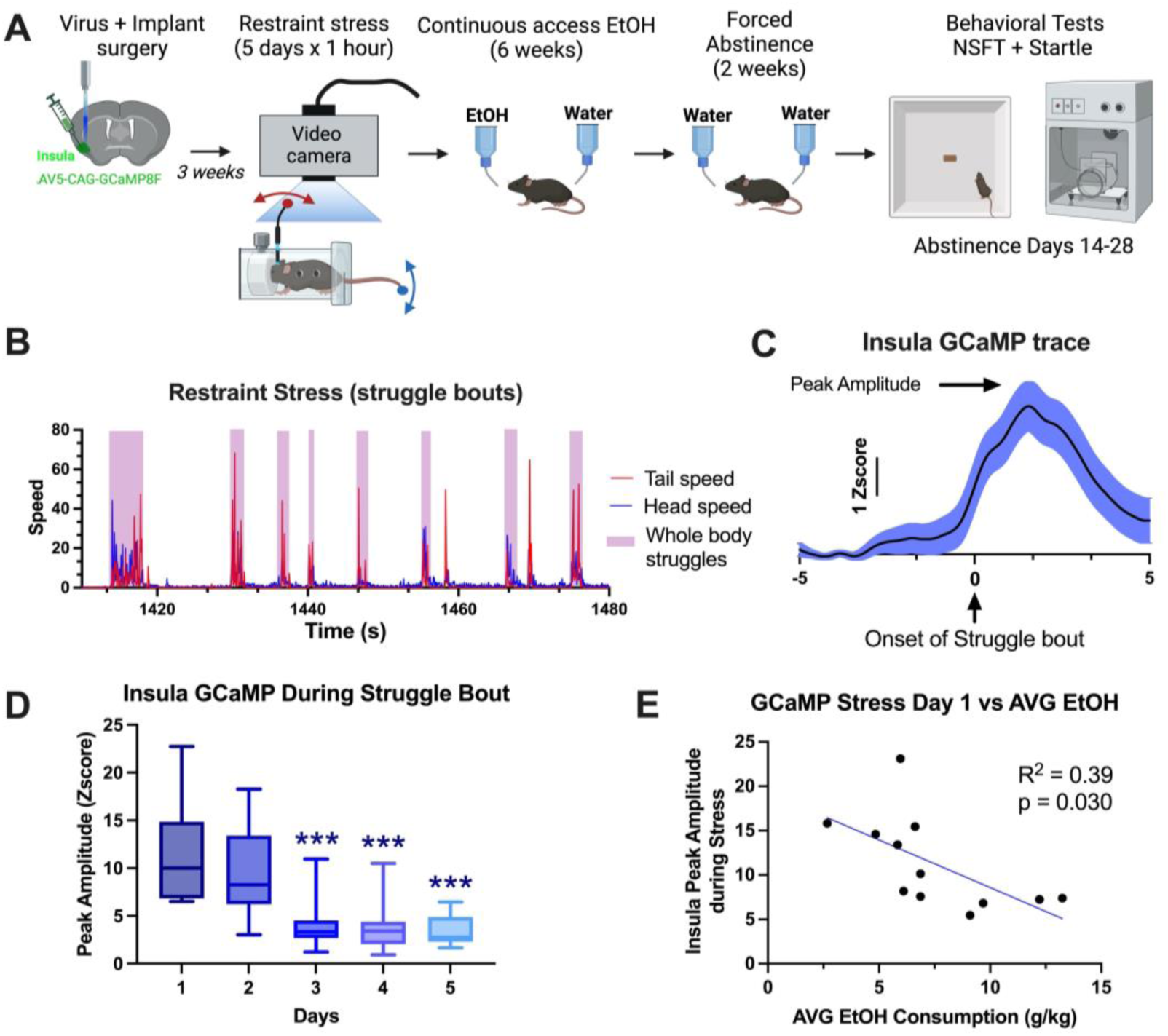
Mid-insular GCaMP response during restraint stress struggle bouts negatively correlates with ethanol drinking in female mice. (A) Experimental design for figures 2 and 3. (B) A representative trace shows the speed of tail, head, and whole-body struggle bouts during restraint stress measured with DeepLabCut and a custom R code. (C) Representative trace of insula GCaMP activity time locked with struggle bouts during restraint stress. (D) Insula GCaMP peak amplitudes time locked to struggle bout onset decrease over 5 days of restraint stress. (E) Peak amplitude of insula GCaMP activity on day 1 of restraint stress negatively correlates with the average ethanol consumption during weeks 4-6. *p<0.05, ***p<0.001

### Mid-insular GCaMP activity during restraint stress and negative affect-like behavior in abstinence

After stress and ethanol drinking, we next tested the relationship between pre-alcohol stress-activated mid-insula activity and negative affect-like behavior during abstinence in the same cohort from Figure 2. Two weeks into abstinence, we measured mid-insula GCaMP activity using fiber photometry during NSFT during each interaction with the food pellet. During NSFT, mice typically approach the food several times before consuming it. The GCaMP signals time-locked with the approach to the food were averaged across subjects, showing time-dependent differences in mid-insula activity (Figure 3A-B). The peak height of the GCaMP signal was higher during the consummatory bout compared to the first approach (paired t-test, t_(12)_=2.81, p=0.0157, Figure 3C). Consistent with the susceptible population identified in Figure 1D, this effect was driven by a subset of mice, outlining a neural correlate to the behaviorally defined population. The peak GCaMP amplitude during the first or last approach did not correlate with the latency to eat (Figure 3D), suggesting the effect was independent of the elapsed time between approaches. A significant positive correlation emerged when comparing mid-insula GCaMP peak amplitude time locked to struggle bout onset during restraint stress with the latency to eat during NSFT (R^2^=0.408, F_(1,8)_=5.52, p=0.0158, Figure 3E). This links heightened stress response to increased anxiety-like behavior in abstinence.

**Figure 3.**
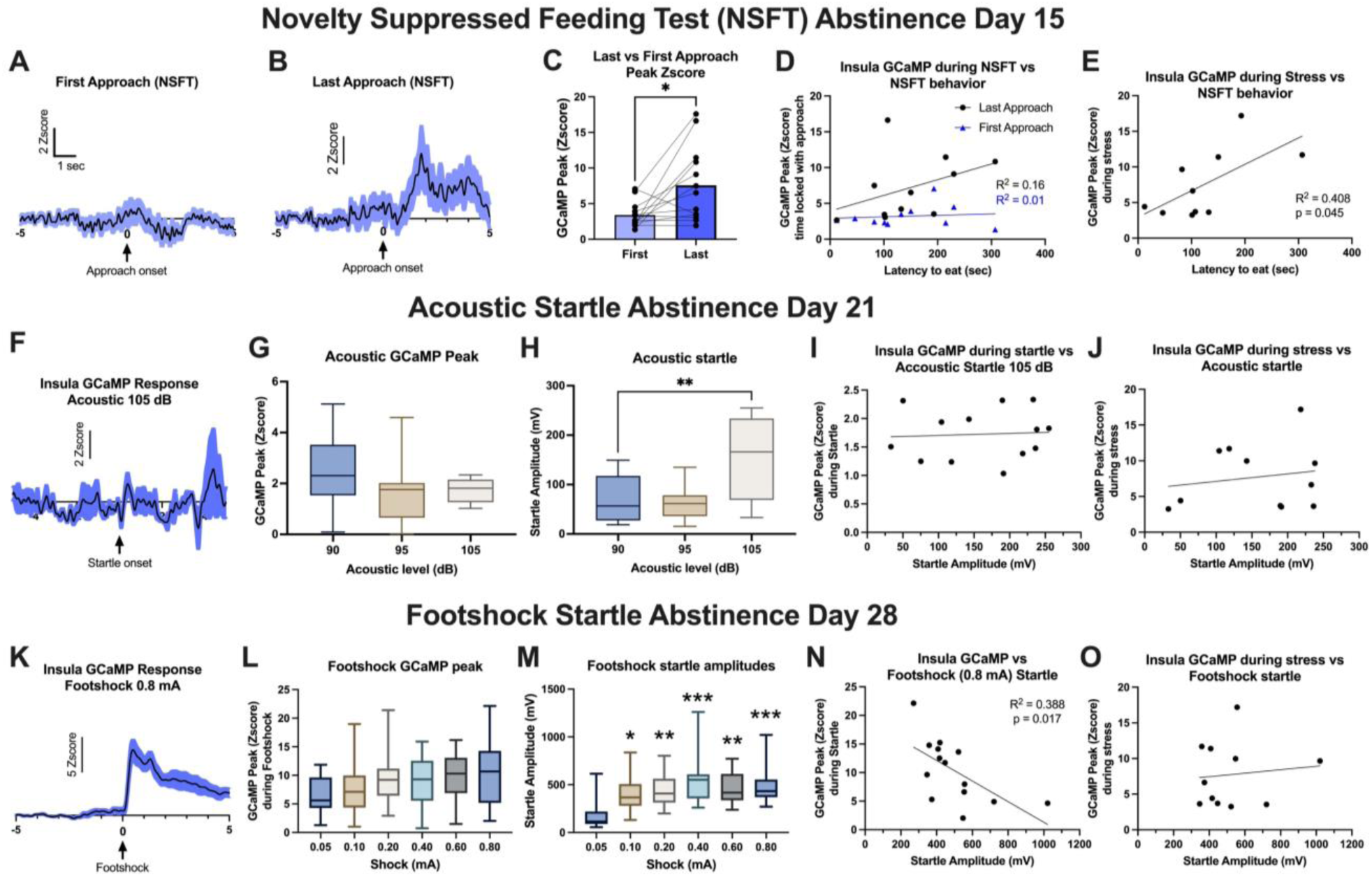
GCaMP activity in the mid-insula is active, and correlates with, negative affect-like behavior during NSFT and footshock startle, but not acoustic startle in female mice. (A-B) Mid-insular GCaMP activity averaged across subjects, time locked with the first and last (consummatory) approaches to the food during NSFT. (C) Mid-insula GCaMP peak amplitude is higher during the last approach compared to the first approach to the food. (D) Insular GCaMP activity during the last approach during NSFT is correlated with the latency to eat during NSFT, but not during the first approach. (E) GCaMP insula activity during restraint stress (eight weeks prior) correlates with latency to eat during NSFT. (F) Representative traces of mid-insula GCaMP activity time locked with 105 dB acoustic startle. (G) Mid-insula GCaMP peak amplitudes during acoustic startle, with no differences across decibels (dB). (H) Startle amplitudes during acoustic startle increase at 105 dB compared to 90 dB. (I-J) Mid-insula GCaMP activity during acoustic startle (I) and restraint stress (J) do not correlate with the acoustic startle amplitude. (K) Representative traces of mid-insula GCaMP activity time locked with 0.8 mA footshock. (L) Mid-insula GCaMP peak amplitudes during footshock startle, with no variation across shock levels. (M) Footshock startle amplitudes increase at higher shock levels. (N) Mid-insula GCaMP activity during footshock startle negatively correlates with the footshock startle amplitude. (O) Mid-insula GCaMP activity during restraint stress eight weeks prior does not correlate with the footshock startle amplitude.*p<0.05, **p<0.01, ***p<0.001

In contrast, mid-insula activity was not recruited during the acoustic startle test on abstinence day 21 (Figure 3F-G), despite increasing decibels evoking a stronger startle response (F_(2,39)_=10.10, p=0.0003, Figure 3H). The behavioral response to the highest decibel (105dB) was not correlated with mid-insula GCaMP activity during stimulus presentation (Figure 3I). Similarly, pre-CDFA restraint stress GCaMP did not correlate with acoustic startle response (Figure 3J).

During the footshock startle test on abstinence day 24, all shock levels elicited a similar amplitude mid-insula GCaMP response (Figure 3K-L). Startle amplitude was higher in the five highest intensities compared to the lowest intensity (0.05mA) (F_(5,78)_=5.789, p=0.0001, Dunnett’s multiple comparisons: 0.05 versus 0.1: p=0.21, 0.05 versus 0.2: p=0.0025, 0.05 versus 0.4: p<0.0001, 0.05 versus 0.6: p=0.0027, 0.05 versus 0.8: p=0.0004, Figure 3M). Comparing each mouse’s peak insula GCaMP amplitude during the highest shock intensity (0.8mA) with the behavioral response revealed a negative correlation between GCaMP and startle response (R^2^=0.388, F_(1,12)_=7.61, p=0.017, Figure 3N), indicating mice with high insula activity during stimulus exposure exhibited a lower behavioral response to the shock stimulus. In contrast, no correlation was observed between pre-CDFA stress activity and footshock startle response (Figure 3O). These results reinforce the unique role of mid-insula activity during pre-alcohol stress in predicting NSFT behavior, but this does not generalize to acoustic or footshock startle.

### The mid-insula to BNST pathway encodes susceptibility to negative affect-like behavior in protracted abstinence

Given the BNST’s innate role in maintaining negative emotional states in abstinence, we investigated the role of the mid-insula-BNST circuit in stress-induced susceptibility to abstinence-induced negative affect with circuit-specific chemogenetics. Female mice received retrograde cre-recombinase injections into the dorsal BNST and cre-dependent Gi-DREADD (hM4Di) into the mid-insula (Figure 4A). Chemogenetically decreasing insula⟶BNST activity with C21 (1mg/kg i.p., 1h before stress) tended to reduce the number of struggle bouts, although this effect was not statistically significant (t_(24)_=1.5, p=0.14, Figure 4C). This manipulation did not impact ethanol consumption during CDFA (Figure 4D).

**Figure 4.**
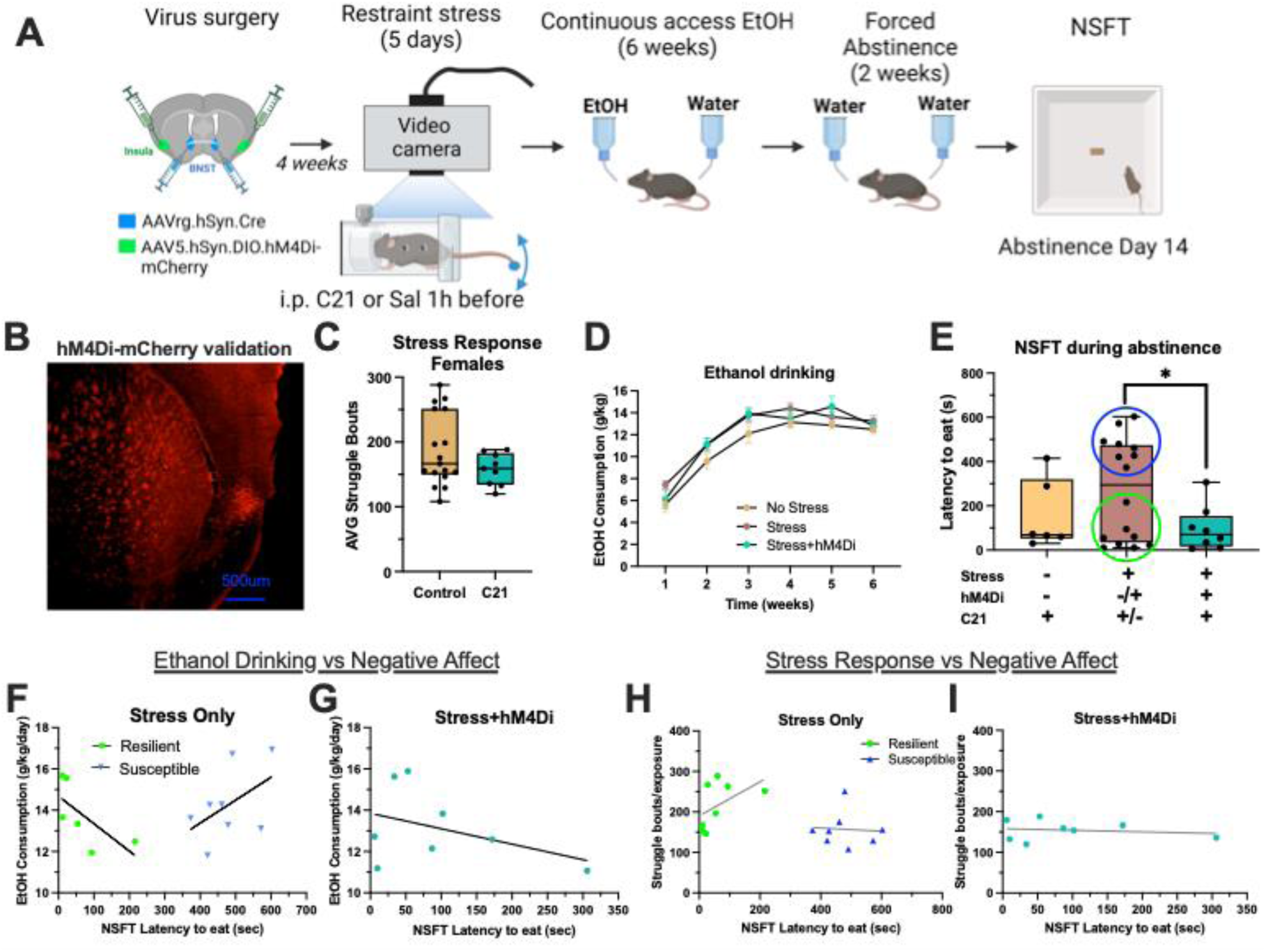
Chemogenetic inhibition of the insula-BNST pathway blocks the emergence of a stress-vulnerable sub-population in female mice. (A) Experimental design. (B) Representative image of hM4Di placement in the mid-insular cortex. (C) The average number of struggle bouts across five days of sub-chronic restraint stress is not different between the control and C21-hM4Di groups. (D) No difference in ethanol drinking levels over six weeks of CDFA between stress, no stress and C21-hM4Di groups. (E) Latency to eat during NSFT in mice after two weeks of abstinence increases in mice with prior stress due to emergence of stress-resilient (green circle) and stress-susceptible (blue circle) sub-populations and is blocked with inhibition of the insula-BNST pathway during prior stress (C21+HM4DI). (F) In stress control mice, average ethanol consumption during weeks 4-6 plotted against the latency to eat during NSFT shows opposing correlation directions in two distinct sub-populations. (G) In insula→BNST hM4Di mice that received C21 during stress, average ethanol consumption shows no correlation with latency to eat during NSFT. (H) The frequency of struggle bouts during restraint stress 8 weeks prior has opposing correlations with latency to eat during NSFT. (I) In insula→BNST hM4Di mice that received C21 during stress, average number of struggle bouts compared against the latency to eat during NSFT. *p<0.05

Two weeks into abstinence, NSFT revealed a similar stress effect, with susceptible and resilient populations emerging. Chemogenetically inhibiting the insula-BNST pathway during stress reduced NSFT (Dunnett’s multiple comparison test: p=0.0471 for stress versus C21 stress, Figure 4E), an effect driven by a lack of a stress-susceptible population. Sub-chronic restraint stress did not produce stress-susceptible and resilient populations in a cohort without subsequent ethanol exposure, as pathway inhibition did not affect NSFT latency (Figure S2). This suggests that stress-resilient and susceptible sub-populations are revealed and later rescued with insula-BNST inhibition only during ethanol abstinence.

Given the larger sample size of susceptible versus resilient populations in the stress + ethanol cohort, we conducted a *post hoc* correlative analysis to compare stress response and drinking patterns with latency to eat during NSFT. This revealed a trend where resilient mice consuming more ethanol showed lower NSFT latency, while susceptible mice with consuming more ethanol exhibited higher latency (Resilient: R^2^=0.46, Susceptible: R^2^=0.23, Figure 4F). Insula-BNST inhibition during pre-ethanol stress also produced a negative trend between drinking and latency (R^2^=0.17, Figure 4G), similar to resilient mice without hM4Di. The number of struggle bouts during pre-CDFA stress did not correlate with resilient, susceptible, or hM4Di groups (Resilient: R^2^=0.25, Susceptible: R^2^=0.004, hM4Di: R^2^=0.02, Figure 4H-I). Thus, despite no direct correlation between struggling during restraint and NSFT latency, chemogenetically inhibiting the insula-BNST pathway during stress disrupts the development of stress-induced susceptibility, leading to lasting behavioral effects into abstinence.

### The mid-insula to BNST pathway and stress-induced susceptibility to negative affect differentially impact male mice

The CDFA model reliably produces negative affect in protracted abstinence in female, but not male mice [35]. To determine whether our newly established model produces similar effects in male mice, we used the same chemogenetic strategy as above. First, we compared male and female struggle behavior during restraint stress independent of circuit manipulation. Female mice exhibited more struggle bouts than male mice (t_(31)_=4.84, p<0.0001, Figure 5A). However, inhibiting insula-to-BNST neurons (C21-hM4Di) during stress prevented the emergence of the sex difference (Figure 5B), driven by a trending increase in male struggle bouts (t_(25)_=1.85, p=0.075, Figure 5C) and a slight decrease in female struggle bouts (Figure 4C).

**Figure 5.**
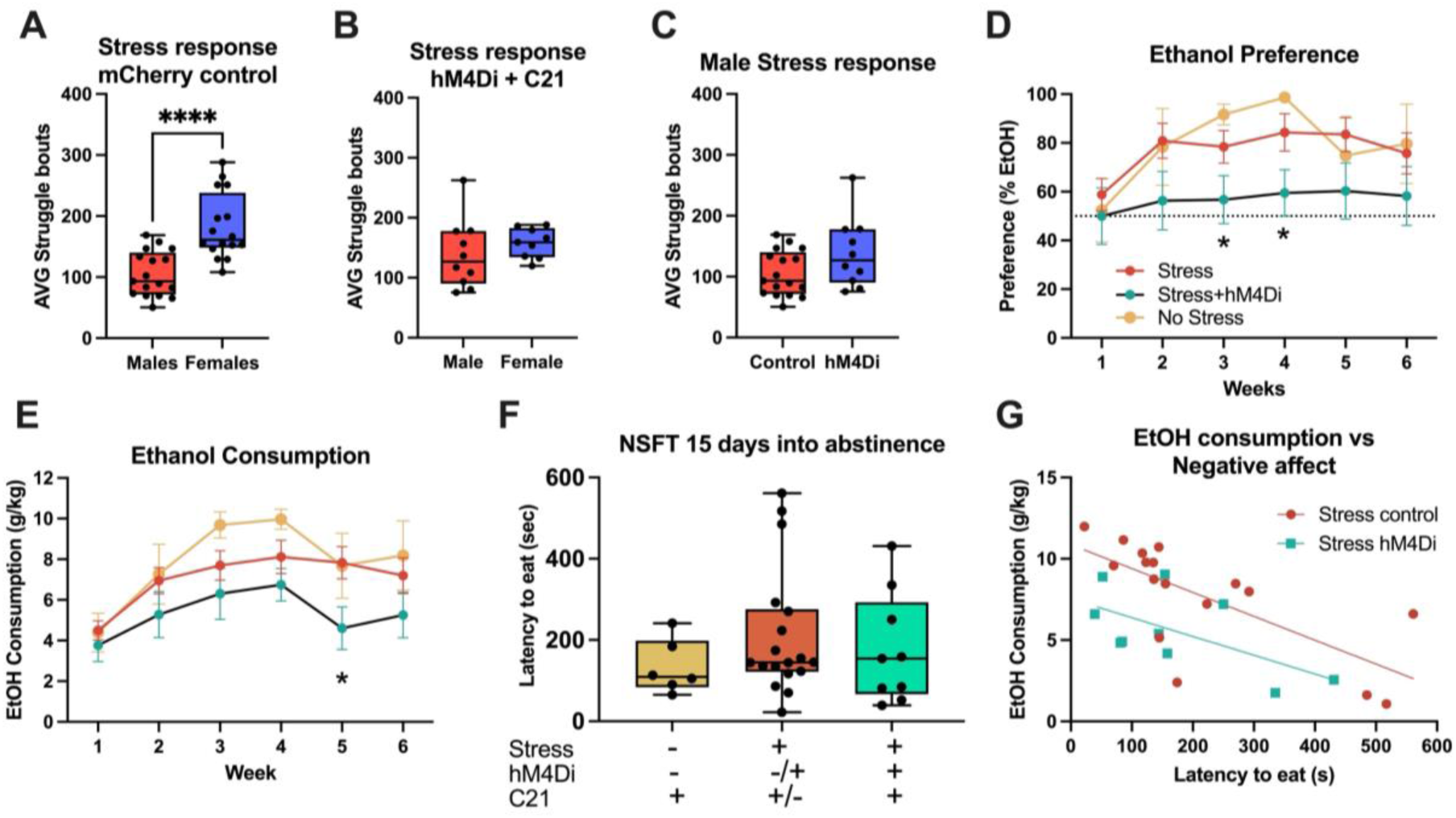
Chemogenetic inhibition of the insula-BNST pathway reduces voluntary ethanol consumption but does not affect negative affective behavior during abstinence in male mice. (A) The average number of struggle bouts during restraint stress is higher in females compared to males (B) No differences in average struggle bouts between males and female mice when the insula-BNST pathway is chemogenetically inhibited during stress (C21+hM4Di). (C) Average number of struggle bouts does not change between control and hM4Di groups in males. (D-E) Ethanol preference over water (D) and consumption (g/kg/day) (E) during CDFA in males are reduced in hM4Di mice compared to stress controls. (F) Latency to eat during NSFT is not affected by prior stress or hM4Di groups in males. (G) EtOH consumption (g/kg/day) negatively correlates with the latency to eat during NSFT across all groups. *p<0.05, ****p<0.0001.

While stress exposure alone did not impact the preference for ethanol over water in male mice, inhibition of the insula-BNST pathway during stress (C21-hM4Di) caused a lower preference for ethanol (Two-Way ANOVA, F_(1,31)_=4.703, p=0.038, Tukey’s post hoc, p<0.05 for weeks 3-4; Figure 5D). A similar effect was evident with consumption. A significant effect of consumption over time (Two-Way ANOVA, F_(3,114)_=15.94, p<0.0001) was driven by the stress+saline and no-stress mice (Tukey’s post hoc p<0.05 for week 1 vs weeks 2-6) and no change in stress+C21 mice (Figure 5E). This same post hoc test revealed a treatment effect at week 5 (saline vs C21 p=0.011). Unlike female mice, stress exposure in males did not impact NSFT latency in abstinence or produce distinct stress-susceptible and resilient mice. C21 similarly did not affect NSFT latency in abstinence (Figure 5F). Male mice showed a significant negative correlation between ethanol consumption (average g/kg/day weeks 4-6) and NSFT latency in abstinence (R^2^=0.51, p=0.002, Figure 5G), with higher consumption associated with lower latency. In hM4Di-C21 male mice, this relationship remained but with a significantly lower y-intercept, but not the slope (F_(1,24)_=9.13, p=0.006, Figure 5G), indicating reduced ethanol consumption compared to controls. These results highlight distinct sex differences in stress-coping behavior, stress-induced drinking patterns, and abstinence-induced negative affect-like behavior, with the insula-BNST playing distinct roles in males and females.

## DISCUSSION

Stress exposure recruits highly individualized behavioral and neural responses. Isolating the unique differences in genetically and environmentally controlled mouse models allows for direct comparisons between a basal stress response and a constant period of heightened negative emotionality, such as abstinence from alcohol in AUD patients. We demonstrate a distinct role for the insula-BNST pathway in encoding basal stress response and susceptibility to hyperkatifeia in protracted abstinence following chronic alcohol drinking. We identified the emergence of two distinct subsets of stress-exposed female C57BL/6J mice-one susceptible population that exhibited heightened negative affect-like behavior in protracted abstinence and one resilient population that resembled unstressed mice in abstinence. Fiber photometry recordings in the mid-insula during restraint stress demonstrate a robust increase in activity time-locked to initiating an active coping behavior (“struggle bout”). The peak amplitude of this time-locked signal correlated with anxiety-like behavior during NSFT, but not startle response tests. Pathway-specific chemogenetic inhibition of the mid-insula neurons projecting to the BNST before each stress exposure prevented the emergence of this stress susceptible population in abstinence. Interestingly, these effects were specific to female mice, as no susceptible population or notable correlations were observed in male mice. Together, we characterized a model for stress and alcohol drinking that uncovers the relationship between stress response and negative affect-like behavior in abstinence in female mice, an understudied population in the treatment of AUD. Embracing the individual variability in stress response and the differential recruitment of stress-sensitive neurocircuits can impact risk assessment, diagnosis, and treatment for stress-related and AUD.

### Stress drives heightened negative affect-like behavior in abstinence in female mice

The primary goal was to engage a quantifiable behavioral response to stress and compare this response to abstinence-induced negative affect, by directly characterizing active neurocircuitry and behavioral response during stress and in abstinence. Sub-chronic restraint stress before CDFA resulted in higher latency to eat during NSFT 15 days into abstinence. However, this effect was not observed in either an acoustic or footshock startle test. A major difference between NSFT and startle tests is the conflict and cognitive processing involved in NSFT, whereas startle tests assess the behavioral response to a sudden, unexpected aversive stimulus. We predicted the startle tests would model a re-exposure to a mild aversive stressor, producing a heightened response in mice with prior stress exposure. While our goal was to test a novel aversive experience, generalization may not occur. Future studies should examine re-exposure to the same stress in abstinence to test this idea. In addition, testing different stress exposures before drinking and the same or different aversive stimuli in abstinence will address the specificity of the stress and abstinence-induced negative affect relationship. Lastly, a startle response reflects an active coping or escape behavior, akin to the struggle bouts quantified during restraint stress. The acoustic startle data for stress only (Figure 1E and 3H-J) was notably more variable than the ethanol-only mice, which may reflect stress-induced variability in passive vs active coping strategies. Shock startle drove a trend towards lower startle response in the stress+CDFA mice, possibly suggesting increased freezing behavior. Testing these trends in a full fear conditioning paradigm may resolve the differences in startle response tests and NSFT.

### Stress reveals susceptibility to heightened negative affect-like behavior in abstinence in female mice

In stress-exposed mice, higher negative affect-like behavior during NSFT in abstinence appeared to be driven by a subset of mice (Figure 1C). The higher powered insula⟶BNST hM4Di experiment (Figure 4) reinforced this, as two similar populations of mice emerged, mice susceptible and resilient to heightened negative affect in abstinence (Figures 1C, 4E). While we predicted this pattern would generalize to other negative affect-like behavioral tests, acoustic or footshock startle response tests did not follow a similar pattern (Figure 1). Tests for hyponeophagia like NSFT can yield bimodal behavior distributions in some models [36], a feature useful for evaluating pharmacological treatments with high non-responder rates, like many antidepressants. Notably, the number of struggle bouts during stress did not correlate with NSFT latency in either resilient or susceptible mice, and the slope of the line was similar between populations. Behavioral output and stress coping strategy do not predict stress susceptibility outright. Instead, the salience and underlying neural response to the stress may better predict susceptibility to negative affect-like behavior in abstinence.

A key finding was the absence of resilient and susceptible populations in male mice (Figure 5). Male mice appeared to be less susceptible to the effects of restraint stress, supported by fewer overall struggle bouts than female mice (Figure 5A). In addition, the CDFA model generally yields lower ethanol consumption and less abstinence-induced negative affect on NSFT in males, highlighting innate sex differences in our model. Overall, this model seems better suited for detecting individual differences in female mice. This emphasizes the importance of developing and utilizing stress and alcohol models that capture resilient and susceptible female phenotypes.

### Sub-chronic stress and continuous access ethanol drinking in C57BL/6J mice

Our approach provides insight into how sub-chronic restraint stress alters drinking behavior in a continuous access model. While the influence of stress on drinking behavior in mice has been widely studied, no consensus has been reached. Several factors influence variability in outcomes, including mouse strain, length and modality of stress, alcohol exposure model, timing of stress relative to alcohol exposure, and sex [37]. In our study, the sub-chronic (five days of one-hour) restraint stress did not impact drinking patterns in female or male mice (Figures 1B, 4D). The absence of a stress effect on ethanol drinking in could relate to the continuous access alcohol model. Alternative models, such as limited access schedules or testing in alcohol-dependent mice, may be better suited to model the binge drinking patterns associated with stress relief or negative reinforcement [38].

The effect of stress on subsequent drinking patterns in female mice has not been widely tested. Human studies suggest females use alcohol to cope with stress at a higher rate than males [39,40]. However, in our model, stress had no impact on alcohol drinking. This disconnect could be attributed to the sub-chronic (5 days) duration, desensitization to the homotypic stressor, or the continuous access drinking model. Future studies should directly compare different stress and alcohol exposure modalities to better understand the nuanced effects of stress on drinking patterns in male and female mice.

### The mid-insula plays distinct, yet overlapping, roles in stress response and abstinence-induced negative affect-like behavior

We link a pre-alcohol stress response in the mid-insula with a behavioral response in abstinence. The strongest relationship was observed with NSFT, where average peak GCaMP amplitude time-locked to struggle bout onset during restraint stress positively correlated with NSFT latency to eat (Figure 3E). Repeated stress exposure reduced time-locked peak GCaMP signal with each exposure, replicating earlier findings [20] and supporting the mid-insula’s role in desensitization after repeated homotypic stressors. Interestingly, mid-insula GCaMP activity during the first food approach was significantly lower than during the last consummatory approach. While this may appear contradictory, we propose a relationship between these signals. During initial restraint stress exposure, a failed escape attempt may signal heightened anxiety.

However, after repeated exposures, the consequence of a failed escape is minimal, as the mouse learns that the stressor will subside regardless of escape success. In NSFT, heightened insula activity during the consummatory approach may reflect increased sensitivity with each approach, culminating in food consumption and a more vulnerable state. The increased insula GCaMP only during the consummatory bout contrasts with prior observations in the dBNST, a downstream target of the mid-insula. dBNST GCaMP signal increased at the onset of all approaches to the food [41], suggesting the mid-insula tips the scale in overcoming hyponeophagia to drive consumption, potentially by modulating interoceptive hunger or shifting food valence through heightened activity relayed to the BNST. Future studies using circuit-specific GCaMP strategies are needed to explore this possibility fully.

### The mid-insula to BNST pathway differentially encodes stress response and alcohol-related behavior in male and female mice

Given insula neurons’ extensive interconnectivity and functional diversity, different populations may be recruited during stress and NSFT in abstinence. For example, a recent study outlines distinct firing patterns in subsets of anterior insula neurons during drinking [42]. We targeted the mid-insula to BNST projection given its role in negative effects following acute stress exposure and during protracted alcohol abstinence [19,20]. The increase in mid-insula GCaMP peak activity during the first versus last approach in NSFT was driven by a subset of mice, suggesting the mid-insula as a neural correlate of the susceptible and resilient populations identified in other experiments. Chemogenetic inhibition of the insula⟶BNST neurons during stress exposure prevented the emergence of a susceptible population in abstinence, establishing this circuit as a critical meditator of future behavioral response during heightened negative emotional states like protracted abstinence.

Manipulating this circuit during stress revealed notable sex differences. Female mice exhibited more struggle bouts during restraint stress than males (Figure 5A), an effect blocked in mice treated with hM4Di. This reduction was driven by lower struggles in female hM4Di mice, while male hM4Di mice showed slightly higher struggles. These findings suggest that the insula→BNST pathway is involved in active versus passive coping strategy, with directionality dependent on sex. Of note, these trends were not predictive of susceptible and resilient female mice in abstinence, as struggle bouts and NSFT latency were not significantly correlated (Figure 4I-J).

Manipulating the insula→BNST pathway during stress impacted drinking in male, but not female mice, trending towards decreased escalation of ethanol preference and consumption in males. While the data are somewhat inconclusive, this could reflect a shift in hedonic value of alcohol, with its aversive properties outweighing the rewarding properties, a phenomenon more pronounced in continuous access compared to limited access models.

### Conclusion

We uncovered a distinct link between stress response and negative affect in protracted alcohol abstinence, revealing unique sex-specific stress effects that define female mice as either susceptible or resilient to abstinence-induced hyperkatifeia. We outline a mechanistic link to the mid-insula, with its projection to the BNST, as having a pivotal role in shaping stress susceptibility. Together, this work underscores the critical need for sex-specific approaches to understanding stress responses and coping strategies. By dissecting how the mid-insula⟶BNST is recruited by stress, we can pave the way for more precise diagnostics and transformative, individualized treatment strategies for AUD.

## AUTHOR CONTRIBUTIONS

Benjamin Williams: Data Curation, Investigation, Software, Formal Analysis, Methodology, Project Administration, Visualization, Writing-Original Draft, Writing-Review & Editing Jincy Little: Investigation, Data Curation, Visualization, Writing-Review & Editing Nathaniel O’Connell: Methodology, Formal Analysis, Data Curation, Visualization, Validation, Writing-Review & Editing Samuel Centanni: Conceptualization, Methodology, Software, Investigation, Resources, Data Curation, Writing-Original Draft, Writing-Review & Editing, Visualization, Supervision, Project Administration, Funding Acquisition.

## REFERENCES

1 Sinha R. Alcohol’s Negative Emotional Side: The Role of Stress Neurobiology in Alcohol Use Disorder. Alcohol Res. 2022;42(1):12.

2 Koob GF, Powell P, White A. Addiction as a Coping Response: Hyperkatifeia, Deaths of Despair, and COVID-19. The American journal of psychiatry. 2020;177(11):1031-37.

3 Sinha R. Stress and substance use disorders: risk, relapse, and treatment outcomes. The Journal of clinical investigation. 2024;134(16).

4 Agabio R, Campesi I, Pisanu C, Gessa GL, Franconi F. Sex differences in substance use disorders: focus on side effects. Addict Biol. 2016;21(5):1030–42.

5 Agabio R, Pisanu C, Gessa GL, Franconi F. Sex Differences in Alcohol Use Disorder. Curr Med Chem. 2017;24(24):2661–70.

6 Bangasser DA, Cuarenta A. Sex differences in anxiety and depression: circuits and mechanisms. Nature reviews Neuroscience. 2021;22(11):674–84.

7 Grossman M, Wood W. Sex differences in intensity of emotional experience: a social role interpretation. J Pers Soc Psychol. 1993;65(5):1010–22.

8 Maciejewski PK, Prigerson HG, Mazure CM. Sex differences in event-related risk for major depression. Psychol Med. 2001;31(4):593–604.

9 Ibrahim C, Rubin-Kahana DS, Pushparaj A, Musiol M, Blumberger DM, Daskalakis ZJ, et al. The Insula: A Brain Stimulation Target for the Treatment of Addiction. Frontiers in pharmacology. 2019;10:720.

10 Menon V, Uddin LQ. Saliency, switching, attention and control: a network model of insula function. Brain Struct Funct. 2010;214(5-6):655–67.

11 Centanni SW, Janes AC, Haggerty DL, Atwood B, Hopf FW. Better living through understanding the insula: Why subregions can make all the difference. Neuropharmacology. 2021;198:108765.

12 Namkung H, Kim SH, Sawa A. The Insula: An Underestimated Brain Area in Clinical Neuroscience, Psychiatry, and Neurology. Trends in neurosciences. 2017;40(4):200–07.

13 Centanni SW, Janes AC, Haggerty DL, Atwood B, Hopf FW. Better living through understanding the insula: Why subregions can make all the difference. Neuropharmacology. 2021:108765.

14 Craig AD. How do you feel--now? The anterior insula and human awareness. Nature reviews Neuroscience. 2009;10(1):59–70.

15 Dager AD, Anderson BM, Rosen R, Khadka S, Sawyer B, Jiantonio-Kelly RE, et al. Functional magnetic resonance imaging (fMRI) response to alcohol pictures predicts subsequent transition to heavy drinking in college students. Addiction. 2014;109(4):585–95.

16 Myrick H, Anton RF, Li X, Henderson S, Drobes D, Voronin K, et al. Differential brain activity in alcoholics and social drinkers to alcohol cues: relationship to craving. Neuropsychopharmacology: official publication of the American College of Neuropsychopharmacology. 2004;29(2):393–402.

17 Bach P, Zaiser J, Zimmermann S, Gessner T, Hoffmann S, Gerhardt S, et al. Stress-Induced Sensitization of Insula Activation Predicts Alcohol Craving and Alcohol Use in Alcohol Use Disorder. Biological psychiatry. 2024;95(3):245–55.

18 Gorka SM, Radoman M, Jimmy J, Kreutzer KA, Manzler C, Culp S. Behavioral and brain reactivity to uncertain stress prospectively predicts binge drinking in youth. Neuropsychopharmacology: official publication of the American College of Neuropsychopharmacology. 2023.

19 Centanni SW, Morris BD, Luchsinger JR, Bedse G, Fetterly TL, Patel S, et al. Endocannabinoid control of the insular-bed nucleus of the stria terminalis circuit regulates negative affective behavior associated with alcohol abstinence. Neuropsychopharmacology: official publication of the American College of Neuropsychopharmacology. 2019;44(3):526–37.

20 Luchsinger JR, Fetterly TL, Williford KM, Salimando GJ, Doyle MA, Maldonado J, et al. Delineation of an insula-BNST circuit engaged by struggling behavior that regulates avoidance in mice. Nature communications. 2021;12(1):3561.

21 Gehrlach DA, Weiand C, Gaitanos TN, Cho E, Klein AS, Hennrich AA, et al. A whole-brain connectivity map of mouse insular cortex. Elife. 2020;9.

22 Koob GF, Zorrilla EP. Neurobiological mechanisms of addiction: focus on corticotropin-releasing factor. Curr Opin Investig Drugs. 2010;11(1):63–71.

23 Reynolds SM, Zahm DS. Specificity in the projections of prefrontal and insular cortex to ventral striatopallidum and the extended amygdala. J Neurosci. 2005;25(50):11757–67.

24 Berry SC, Wise RG, Lawrence AD, Lancaster TM. Extended-amygdala intrinsic functional connectivity networks: A population study. Hum Brain Mapp. 2021;42(6):1594–616.

25 Flook EA, Feola B, Avery SN, Winder DG, Woodward ND, Heckers S, et al. BNST-insula structural connectivity in humans. NeuroImage. 2020;210.

26 Marino RAM, Girven KS, Figueiredo A, Navarrete J, Doty C, Sparta DR. Binge ethanol drinking associated with sex-dependent plasticity of neurons in the insula that project to the bed nucleus of the stria terminalis. Neuropharmacology. 2021;196:108695.

27 Guidelines for the Care and Use of Mammals in Neuroscience and Behavioral Research. Washington (DC); 2003.

28 Holleran KM, Wilson HH, Fetterly TL, Bluett RJ, Centanni SW, Gilfarb RA, et al. Ketamine and MAG Lipase Inhibitor-Dependent Reversal of Evolving Depressive-Like Behavior During Forced Abstinence From Alcohol Drinking. Neuropsychopharmacology: official publication of the American College of Neuropsychopharmacology. 2016;41(8):2062–71.

29 Holleran KM, Wilson HH, Fetterly TL, Bluett RJ, Centanni SW, Gilfarb RA, et al. Ketamine and MAG Lipase Inhibitor-Dependent Reversal of Evolving Depressive-Like Behavior During Forced Abstinence From Alcohol Drinking. Neuropsychopharmacology. 2016.

30 Olsen CM, Winder DG. Operant sensation seeking in the mouse. J Vis Exp. 2010(45).

31 Pang TY, Renoir T, Du X, Lawrence AJ, Hannan AJ. Depression-related behaviours displayed by female C57BL/6J mice during abstinence from chronic ethanol consumption are rescued by wheel-running. The European journal of neuroscience. 2013;37(11):1803–10.

32 Centanni SW, Conley SY, Luchsinger JR, Lantier L, Winder DG. The impact of intermittent exercise on mouse ethanol drinking and abstinence-associated affective behavior and physiology. Alcoholism, clinical and experimental research. 2022;46(1):114–28.

33 Grissom N, Kerr W, Bhatnagar S. Struggling behavior during restraint is regulated by stress experience. Behavioural brain research. 2008;191(2):219–26.

34 Patel S, Roelke CT, Rademacher DJ, Hillard CJ. Inhibition of restraint stress-induced neural and behavioural activation by endogenous cannabinoid signalling. The European journal of neuroscience. 2005;21(4):1057–69.

35 Salazar AL, Centanni SW. Sex Differences in Mouse Models of Voluntary Alcohol Drinking and Abstinence-Induced Negative Emotion. Alcohol (Fayetteville, NY). 2024;121:45–57.

36 Samuels BA, Leonardo ED, Gadient R, Williams A, Zhou J, David DJ, et al. Modeling treatment-resistant depression. Neuropharmacology. 2011;61(3):408–13.

37 Becker HC, Lopez MF, Doremus-Fitzwater TL. Effects of stress on alcohol drinking: a review of animal studies. Psychopharmacology. 2011;218(1):131–56.

38 Cho SB, Su J, Kuo SI, Bucholz KK, Chan G, Edenberg HJ, et al. Positive and negative reinforcement are differentially associated with alcohol consumption as a function of alcohol dependence. Psychol Addict Behav. 2019;33(1):58–68.

39 Fox HC, Sinha R. Sex differences in drug-related stress-system changes: implications for treatment in substance-abusing women. Harvard review of psychiatry. 2009;17(2):103–19.

40 Guinle MIB, Sinha R. The Role of Stress, Trauma, and Negative Affect in Alcohol Misuse and Alcohol Use Disorder in Women. Alcohol Res. 2020;40(2):05.

41 Jaramillo AA, Williford KM, Marshall C, Winder DG, Centanni SW. BNST transient activity associates with approach behavior in a stressful environment and is modulated by the parabrachial nucleus. Neurobiology of Stress. 2020;13:100247.

42 Starski P, Morningstar MD, Katner SN, Frasier RM, De Oliveira Sergio T, Wean S, et al. Neural Activity in the Anterior Insula at Drinking Onset and Licking Relates to Compulsion-Like Alcohol Consumption. The Journal of neuroscience: the official journal of the Society for Neuroscience. 2024;44(9).

