## Supplemental Material for "A stress-activated mid-insula to BNST pathway regulates susceptibility to abstinence-induced negative affect in female mice"

#### Supplemental Methods

*Animals*

Male and Female C57BL/6J mice (n=124; The Jackson Laboratory; Bar Harbor, ME) were delivered at seven weeks of age, acclimated in standard group housing for one week, and then singly housed and acclimated for one week prior to stress exposure. All mice were maintained on a 12h light/dark cycle (lights on at 0600 hours) under controlled temperature (20-26°C) and humidity (30-70%) levels. Mice were given access to food and water *ad libitum*. All experimental procedures were conducted with the approval of the Institutional Animal Care and Use Committee at Wake Forest University and were within the guidelines set forth by the Care and Use of Mammals in Neuroscience and Behavioral Research [1].

*Stereotaxic Surgeries*

Mice (>8 weeks old) were anesthetized with isoflurane (3% initial dose, 1.5% maintenance dose) for intracranial recombinant adenosine-associated virus (AAV) injection and fiberoptic cannula implant surgeries, using a Leica Angle Two Small Animal Stereotaxic Instrument. The viruses AAV1-hSyn-jGCaMP8f-WPRE, AAV5-hSyn-DIO-hM4Di, and AAV5-hSyn-Cre were used. Mice were injected with 300 nL of the indicated AAV at a rate of 50 nL/min driven by a Micro4 MicroSyringe pump (World Precision Instruments) into the specified region. The needle (World Precision Instruments Nanofil syringe fitted with a Nanofil 33 G Blunt Needle) remained in place for an additional 5 min. A minimum period of 3 weeks prior to further experimentation allowed for full viral expression and recovery. Injection and implantation sites included the mid-insula (from Bregma: AP = 0.02, ML ± 3.66, DV = −4.30, 0° angle) and dorsal BNST (from Bregma: AP =0.20, right ML -0.63, left ML -2.13, DV = −3.88, 15.0° angle). All mice received 5 mg/kg injections of meloxicam once a day for 2 days following surgery.

Chronic optical fiber implantation procedures.

Once exposed, the skull was etched with a gel etchant (Kerr Dental). One stainless-steel mounting screw (PlasticsOne) was installed in the ipsilateral parietal plate posterior to the implant hole. Once the implant was inserted into the brain, it was bonded by applying Optibond primer followed by Optibond adhesive, which was then cured with UV light. Herculite enamel was molded around the screw, implant, and skull. The enamel was then cured with UV light. Mono fiberoptic cannula (Doric Lenses) were tested prior to implantation to ensure >90% power output.

*Restraint Stress*

Restraint stress was conducted as previously described [2]. In brief, following a one-hour acclimation period and 30 minutes after drug administration when applicable, mice were placed into a custom restraining device, RESTRAINT [2,3], for one hour. Videos were recorded using a custom data acquisition pipeline [4] and subsequently run through the DeepLabCut pose estimation pipeline [5]. A custom R script was used to determine the independent velocity of body movement while in the tube. A whole-body struggle bout was quantified as simultaneous head and tail tip movement.

*Chronic Drinking Forced Abstinence Model*

Chronic drinking followed by forced abstinence was done as described previously [6]. Briefly, beginning at 7–8 weeks of age, mice were single-housed and given access to two sipper bottles filled with water. After 1 week of habituation, our experimental group was given access to increasing ethanol concentrations (3% for 3 days, 7% for 7 days) until reaching a maintenance concentration of 10% for the remaining 5 weeks. The control group continued to drink from two bottles that both contained water for the duration of the experiment. After 6 weeks of ethanol or water, ethanol was removed, commencing a forced abstinence period where behavioral testing took place.

*Abstinence Behavioral Tests*

Behavioral assessments of affective state were conducted as follows:

<u>Novelty Suppressed Feeding Test</u>

The novelty suppressed feeding test (NSFT) was conducted as previously described [6-9]. In brief, 13 days into abstinence, mice were food-restricted for the 48 hours leading up to the test. Food access was granted for a 2-hour period, 23-25 hours prior to NSFT. On day 15 of abstinence, mice were subjected to NSFT. Mice were placed into an open arena (50cm x 50cm) with a food pellet in the center of the brightly lit apparatus (300 lux). Latency to eat was measured as the amount of time elapsed before taking a bite of the food pellet. The latency to eat was assessed in real time by the experimenter and confirmed through video recording after the experiment. Mice were removed immediately after the first bite and placed back into their home cage with a pre-weighed food pellet. After 10 minutes the food pellet was reweighed to determine home cage consumption.

Acoustic Startle Response

Testing was conducted in sound-attenuated startle chambers (SRLAB, San Diego Instruments, San Diego, CA), each consisting of a Plexiglas cylinder (3.8 cm diameter) mounted on a Plexiglas platform, with a high-frequency loudspeaker (28 cm above the cylinder) producing all acoustic stimuli. The background noise of each chamber was 70 dB. Movements within the cylinder were detected and transduced by a piezoelectric accelerometer and then digitized and stored by a PC-type computer. Three different acoustic startle levels were presented for each subject: startle pulse of 105 dB, 95 dB, or 90 dB, with three trials given for each startle level. Trials were given in random order. The average inter-trial interval was 67 seconds. Startle amplitudes were averaged across the different stimulus trials for each mouse prior to statistical analyses. Elevated startle amplitudes were interpreted as anxiety-like behavior.

Foot Shock Startle Response

Testing was conducted in the same chamber as previously described in the acoustic startle response. Footshock voltage levels were (mV) 0.05, 0.1, 0.2, 0.4, 0.6 and 0.8, with three trials for each shock level. Trials were randomized similarly to the acoustic startle.

*Fiber Photometry*

Fiber photometry was recorded using a TDT RZ10X fiber photometry system with Synapse acquisition software. Briefly, light from the 470 nm, 17.2mW (Min) fiber-coupled LED (Thorlabs) and light from the 405 nm, 19.3mW (Min) fiber-coupled LED (Thorlabs) was directed into a fluorescence mini cube with six ports and a built-in detector head (Doric Lenses), with spectral bandwidths of 405 and 470 nm. A 405 nm light was modulated at 217 Hz, while 470 nm light was modulated at 330 Hz. Power output was maintained at 20 mA with a DC offset of 3mA for both wavelengths. The light was then directed through a low-autofluorescence mono fiberoptic patch cord with a 400 µm core (Doric). This fiber was connected to the mono fiberoptic cannulas that were implanted into the insula (4mm length, 400/430 I/OD, 0.66 NA, 2.5mm diameter). The power output at the fiber tip was 25–30 µW. Fluorescent emission from the tissue was collected through the same fiber and was detected using a femtowatt photoreceiver. Signal acquisition was 1 kHz and low-pass filtered at 6 Hz. The 405 nm excitation channel served as an isosbestic, calcium-independent control wavelength for GCaMP, allowing for bleaching and movement artifact corrections when directly fit to the calcium-dependent 470 nm channel. MATLAB scripts from TDT were used to fit the 405 nm signal to the 470 nm signal using linear regression. Change in GCaMP mediated signal was calculated as

$$\frac{\Delta F}{F} = \frac{change\ in\ 470\ nm\ signal\ change\ in\ 405nm\ signal}{change\ in\ 405\ nm\ induced\ signal} \quad (1)$$

Time-locked Z scores were then calculated from GCaMP-mediated signals as:

$$Z = \frac{\frac{\frac{\Delta F}{F}}{mean\ of\ \frac{\Delta F}{F}}}{standard\ deviation\ of\ \frac{\Delta F}{F}} \quad (2)$$

*Statistics*

Correlation analysis was conducted using a Pearson's correlation coefficients reported as r with [95% confidence interval]. P-value for correlation analysis was determined with the slope of the line was significantly different from a zero-slope. NSFT analysis was conducted using a parametric two-tailed unpaired Student's *t*-test. Data were analyzed using GraphPad Prism 10. The Machine learning algorithm DeepLabCut was used in conjunction with custom-written R code to quantify struggle bout behaviors during restraint stress, and for in-depth analysis of mouse behavior during NSFT, which includes the number of approaches to the food and the interaction time with the food before taking a bite. For continuous outcomes (e.g., ethanol consumption) across multiple treatment/comparative groups, ANOVA was used with an appropriate method for multiple comparisons (e.g., Dunnet's test) as relevant to the hypotheses being tested. For simple two-group comparisons, t-tests were used.

### References


1 Guidelines for the Care and Use of Mammals in Neuroscience and Behavioral Research. Washington (DC); 2003.

2 Luchsinger JR, Fetterly TL, Williford KM, Salimando GJ, Doyle MA, Maldonado J, et al. Delineation of an insula-BNST circuit engaged by struggling behavior that regulates avoidance in mice. Nature communications. 2021;12(1):3561.

3 Williford KM, Taylor A, Melchior JR, Yoon HJ, Sale E, Negasi MD, et al. BNST PKCdelta neurons are activated by specific aversive conditions to promote anxiety-like behavior. Neuropsychopharmacology : official publication of the American College of Neuropsychopharmacology. 2023.

4 Centanni SW, Smith ACW. PiRATeMC: A highly flexible, scalable, and low-cost system for obtaining high quality video recordings for behavioral neuroscience. Addict Neurosci. 2023;8.

5 Mathis A, Mamidanna P, Cury KM, Abe T, Murthy VN, Mathis MW, et al. DeepLabCut: markerless pose estimation of user-defined body parts with deep learning. Nature neuroscience. 2018;21(9):1281-89.

6 Centanni SW, Morris BD, Luchsinger JR, Bedse G, Fetterly TL, Patel S, et al. Endocannabinoid control of the insular-bed nucleus of the stria terminalis circuit regulates negative affective behavior associated with alcohol abstinence. Neuropsychopharmacology : official publication of the American College of Neuropsychopharmacology. 2019;44(3):526-37.

7 Holleran KM, Wilson HH, Fetterly TL, Bluett RJ, Centanni SW, Gilfarb RA, et al. Ketamine and MAG Lipase Inhibitor-Dependent Reversal of Evolving Depressive-Like Behavior During Forced Abstinence From Alcohol Drinking. Neuropsychopharmacology. 2016.

8 Pang TY, Renoir T, Du X, Lawrence AJ, Hannan AJ. Depression-related behaviours displayed by female C57BL/6J mice during abstinence from chronic ethanol consumption are rescued by wheel-running. The European journal of neuroscience. 2013;37(11):1803-10.

9 Vranjkovic O, Winkler G, Winder DG. Ketamine administration during a critical period after forced ethanol abstinence inhibits the development of time-dependent affective disturbances. Neuropsychopharmacology : official publication of the American College of Neuropsychopharmacology. 2018.

### Supplemental Figures

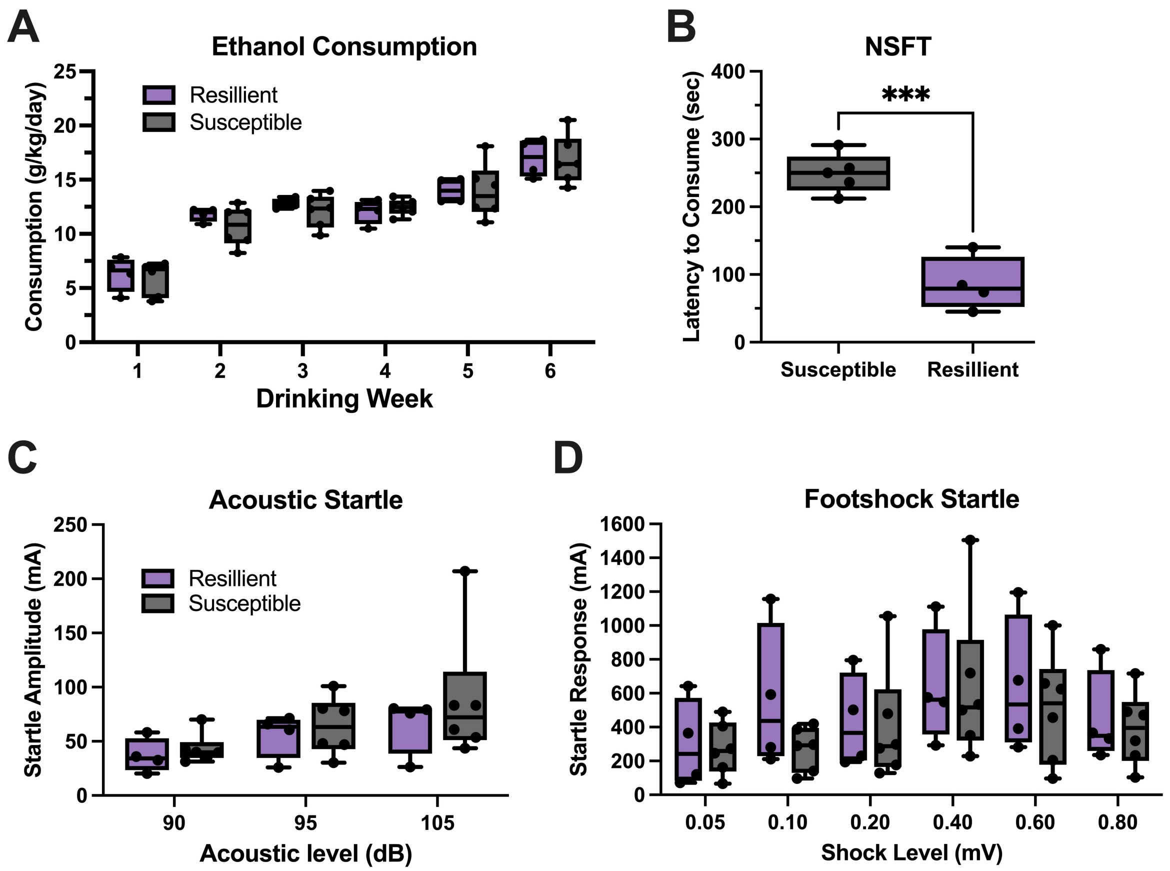


**Figure S1. Negative affective behavior during protracted abstinence between susceptible and resilient stress populations** (A) Ethanol consumption after sub-chronic restraint stress is similar between populations. (B) Latency to eat the food during NSFT in abstinence is significantly lower in resilient mice. (C) Acoustic startle response is similar between susceptible and resilient mice. (D) Foot shock startle response is not different between susceptible and resilient mice. ***$p<0.001$

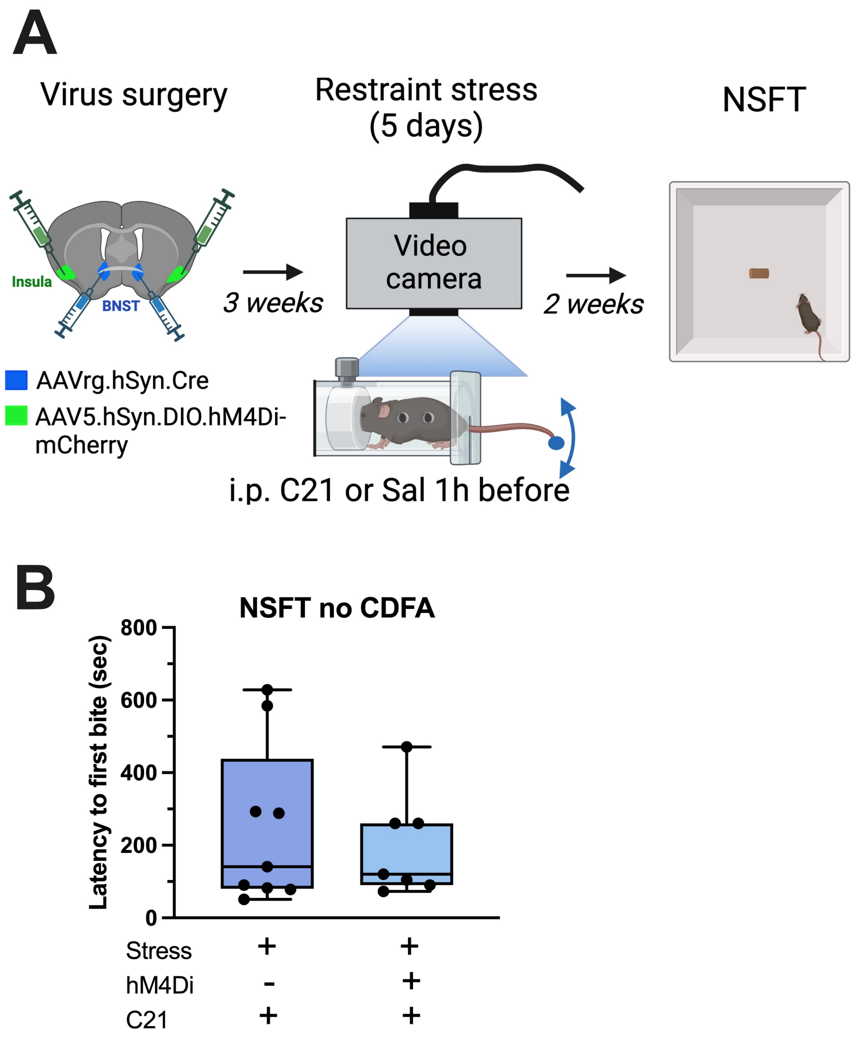


**Figure S2. Inhibition of the insula-BNST pathway during stress has no effect on negative affect in ethanol naïve female mice.** (A) Experiment design. (B) Novelty-suppressed feeding test (NSFT) after 15 days following sub-chronic restraint stress. Inhibiting insula→BNST neurons with hM4Di+C21 does not impact NSFT latency 15 days after sub-chronic stress.
